# miR-203 drives breast cancer cell differentiation

**DOI:** 10.1101/2023.01.23.525208

**Authors:** Nuria G. Martínez-Illescas, Silvia Leal, Patricia González, Osvaldo Graña-Castro, Juan José Muñoz-Oliveira, Alfonso Cortés-Peña, Miguel Quintela-Fandino, Eva Ciruelos, Consuelo Sanz, Sofía Aragón, Leisy Sotolongo, Sara Jiménez, Eduardo J. Caleiras, Francisca Mulero, Cristina Sánchez, Marcos Malumbres, María Salazar-Roa

## Abstract

A hallmark of many malignant tumors is dedifferentiated (immature) cells bearing slight or no resemblance to the normal cells from which the cancer originated. Tumor dedifferentiated cells exhibit a higher capacity to survive to chemo and radiotherapies and have the ability to incite tumor relapse. Inducing cancer cell differentiation would abolish their self-renewal and invasive capacity and could be combined with the current standard of care, especially in poorly differentiated and aggressive tumors (with worst prognosis). However, differentiation therapy is still in its early stages and the intrinsic complexity of solid tumor heterogeneity demands innovative approaches in order to be efficiently translated into the clinic. We demonstrate here that microRNA 203, a potent driver of differentiation in pluripotent stem cells (ESCs and iPSCs), promotes the differentiation of mammary gland tumor cells. Combining mouse *in vivo* approaches and both mouse and human-derived tridimensional organoid cultures, we report here that miR-203 influences the self-renewal capacity, plasticity and differentiation potential of breast cancer cells, and prevents tumor cell growth in vivo. Our work sheds light on differentiation-based antitumor therapies and offers miR-203 as a promising tool for directly confronting the tumor-maintaining and regeneration capability of cancer cells.

## Introduction

Solid tumors are heterogeneous in their cell composition, with a small subset of tumor cells sharing certain biological and molecular properties with tissue-specific stem cells (SC) (1–3). This cell population has long-term renewal potential, supporting a proliferation hierarchy among cancer cells. In some tumors, the genes and signaling pathways that regulate normal SC roles also function as oncogenes or regulate tumor maintenance and progression (4–16). One of the best-characterized examples is WNT–β-catenin signaling, which is essential for the maintenance and proliferation of SCs (17). Importantly, this pathway is frequently mutated in colorectal cancers (18) and is required to sustain tumor growth and progression in several different types of cancers, including colorectal cancer, leukemia and skin basal cell carcinoma (19). Consistent with the reprogramming of tumor cells into an embryonic-like fate, similarities between embryonic mammary SCs and the basal-like and HER2-positive breast cancer subtypes (which are less differentiated than other breast cancer subtypes) have also been described (20). All those observations are consistent with the idea that, for tumor initiation, adult cells are required to undergo reprogramming to a progenitor-like fate.

Many of the current therapeutic strategies aimed at eliminating cancer cells involve treatment with standard anti-proliferative chemotherapy, which often has limited benefits. The residual population of chemotherapy-resistant tumor cells capable of regenerating the disease is- at least by definition-enriched in cancer stem cells (CSCs) (1). This fact has inspired the design of numerous anti-tumor therapies directly targeting the CSC niche, based on inducing their terminal differentiation. Indeed, the original idea of anti-CSC therapy arose in the 1970s and 1980s, from the observation that leukemic cells are blocked in an undifferentiated state. The use of all-trans retinoic acid induced terminal differentiation of leukemic cells (21)—and currently is the standard of care for the treatment of patients with acute promyelocytic leukemia. The success of all-trans retinoic acid therapy inspired other therapies that were based on inhibiting epigenetic regulators to induce cancer differentiation in multiple hematological malignancies (22) and the same mechanism also shows certain promise in solid tumors. Nevertheless, it is now established that even differentiated cells can be reprogrammed into stem-like cells, suggesting that cell state reprogramming is more common and occurs in more diverse cell types than previously thought (23, 24). Indeed, this type of reprogramming can be used to re-establish stem-like hierarchies in tumors even after elimination of putative CSCs (25). Therefore, eliminating unstable cells and also abrogating the mechanisms by which tumor cells gain cell state plasticity may be the most productive differentiation strategy. Such complexity makes the enticing therapeutic targeting of undifferentiated cancer cells still uncertain.

Novel players in carcinogenesis are microRNAs (miRNAs), which are epigenetically regulated but also control epigenetic events (26). miRNAs comprise a class of small noncoding RNAs that is involved in posttranscriptional regulation of gene expression. miRNAs act by inhibiting translation of target mRNAs, and it is estimated that one-third of protein-coding mRNAs are subjected to regulation by miRNA. miRNA deregulation has been implicated in cancer development, and both oncogenic and tumor-suppressor miRNAs have been identified, many of which act through inhibition of translation of proteins controlling cell proliferation, survival, and development (26).

Fuchs’ laboratory first described an *in vivo* role for microRNA 203 as a suppressor of stemness in developing epidermis (27). Soon after that, we described miR-203 as a tumor suppressor in hematopoietic tumors. Our lab found that miR-203 expression was frequently silenced in mouse and human T and B cell malignancies through hypermethylation of its genomic region, and ABL1 and BCR-ABL1 fusion transcripts are indeed direct targets of miR-203-mediated translational repression (28). The same year, a landmark paper from Massague’s group identified a set of eight microRNAs whose expression was inversely correlated with the metastatic potential of human breast cancer cell lines (29). Though not studied further, miR-203 was among the eight miRNAs initially identified in that study. In the years since this report was published, miR-203 has been shown to regulate genes involved in crucial tumor pathways, such as signal transduction (BCR-ABL1), stemness (p63, BMI1), migration (LASP1, ASAP1), as well as known regulators of metastasis (SNAI1/2) among many others (30–39). However, the capacity of miR-203 to fine-tune the cancer cell differentiation remains uncertain, and deserves a more focused research.

Recently, our laboratory has identified an unprecedented role of miR-203 modulating both reprogramming from somatic to pluripotent cells (40) and the differentiation capacity of stem cells (41). Our data support the intriguing fact that a brief exposure to miR-203 blocks reprogramming to pluripotency while expanding the differentiation efficiency of stem cells. Such effects are mediated by direct or indirect targeting of the epigenetic landscape, making pluripotent cells more proficient for subsequent differentiation.

Given the obvious parallelisms between tumorigenesis and pluripotency (42–44), we evaluated the outcomes of miR-203 treatment on cancer cell differentiation. Using the classical MMTV-PyMT transgenic mice as a breast cancer model (45), we demonstrate here that miR-203-mediated effects on cellular reprogramming and cell differentiation can be advantageous in antitumor therapy. Combining *in vivo* approaches and their direct version on *in vitro* settings by tumor-derived organoids, we show that a brief exposure to miR-203 controls the self-renewal and proliferative capacity of breast cancer cells, attenuates migratory abilities and provokes a switch from a basal tumor phenotype to a more differentiated luminal-like status, similar to that observed in non-tumor cells.

## Results

### Different schedules of miR-203 treatment prevent tumor initiation, growth and metastasis in the MMTV-PyMT breast cancer mouse model

To easily manipulate miR-203 levels *in vitro* and *in vivo*, we generated a tetracycline-inducible knock-in model in which the miR-203-encoding sequence was inserted downstream of the type I collagen gene and expressed under the control of a tetracycline-responsive element [*ColA1* (miR-203) allele] in the presence of tetracycline reverse transactivator, expressed from the Rosa26 locus [Rosa26 (rtTA) allele] (41). The treatment of *ColA1 (miR-203/miR-203); Rosa26 (rtTA/rtTA)*-derived cells with doxycycline (Dox) leads to a significant induction of miR-203 levels, only when exposed to Dox treatment (41).

To illustrate the role of miR-203 as an antitumor agent *in vivo*, we chose the PyMT breast cancer model for its close similarity to human breast cancer, exemplified by the fact that in these mice a gradual loss of steroid hormone receptors (estrogen and progesterone) and β1-integrin is associated with overexpression of ERBB2 and cyclin D1 in late-stage metastatic cancer (46). In the Tg(MMTV-PyVT) model (also known as “MMTV-PyMT”), transgenic mice express the Polyoma Virus middle T (PyMT) antigen under the direction of the mouse mammary tumor virus (MMTV) promoter/enhancer (47). Hemizygous MMTV-PyMT females develop palpable mammary tumors that metastasize to the lung, and exhibit high penetrance and early onset of mammary cancer compared to other mammary tumor models. Tumor formation and progression in this murine model is notably similar to that observed in patients and is characterized by four stages: hyperplasia, adenoma/mammary intra-epithelial neoplasia, early carcinoma and late carcinoma. Therefore, we crossed our miR-203 inducible mice with the Tg(MMTV-PyVT) model, in order to generate an *in vivo* tool where easily fine-tune the miR-203 levels by Dox treatment in diet, at different time points during mammary tumor development.

We dissected the *in vivo* antitumor effects of miR-203 by inducing its expression at different schedules: (i) starting at tumor onset and sustaining the treatment during two weeks (Fig. 1); (ii) starting at tumor onset and sustaining the treatment throughout the experiment, to the human experimental endpoint (Fig. 2) or (iii) starting once the tumors are established and under exponential growth, and treating every two weeks (Fig. 3). Tumors were followed by micro-CT and the tumor volume was determined. The potential effects of miR-203 on metastasis incidence in the lungs were also evaluated by micro-CT throughout the three *in vivo* experiments, and by histopathology analysis of lung samples at the endpoint in all conditions tested.

**Fig. 1.**
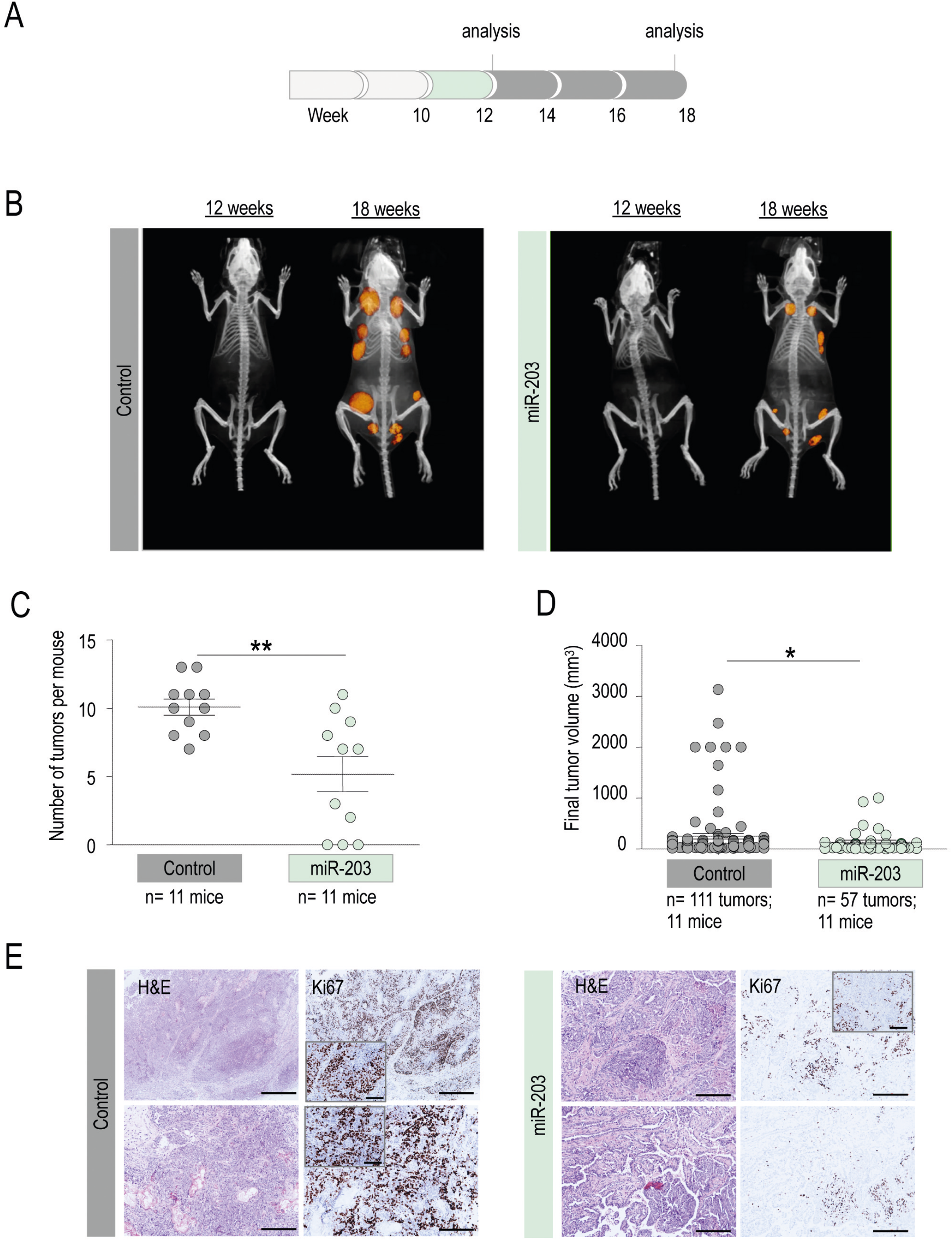
*In vivo* effects of miR-203 treatment on PyMT mice, started at tumor onset and sustained for two weeks. A, Schematic of the Doxycycline (Dox) treatment (in green) schedule *in vivo*, on *miR-203 wild-type* or *miR-203 knock-in*; *PyMT* mice, during two weeks from tumor onset (before the tumors are detected by micro-CT). B, Representative micro-CT images of mice subjected to Dox treatment (in the figures, “control” indicates miR-203 wild-type; “miR-203” indicates knock-in mice), after Dox treatment (12 weeks of age) and at the endpoint (18 weeks of age). C, Number of tumors per mouse at the endpoint, in control and miR-203-treated mice. D, Final tumor volume of control and miR-203-treated mice. In (C, D), data are represented as mean ± s.d. (number of mice and total number of tumors per group are indicated in the figure). *p<0.05; **<0.01 (Student’s t-test). E, Illustrative hematoxylin and eosin (H&E) and Ki-67 immunohistochemistry (IHC) staining of control and miR-203-treated tumors at the endpoint. Two representative tumors are shown per condition. Insets show a higher magnification for details. Scale bar, 500 μm.

**Fig. 2.**
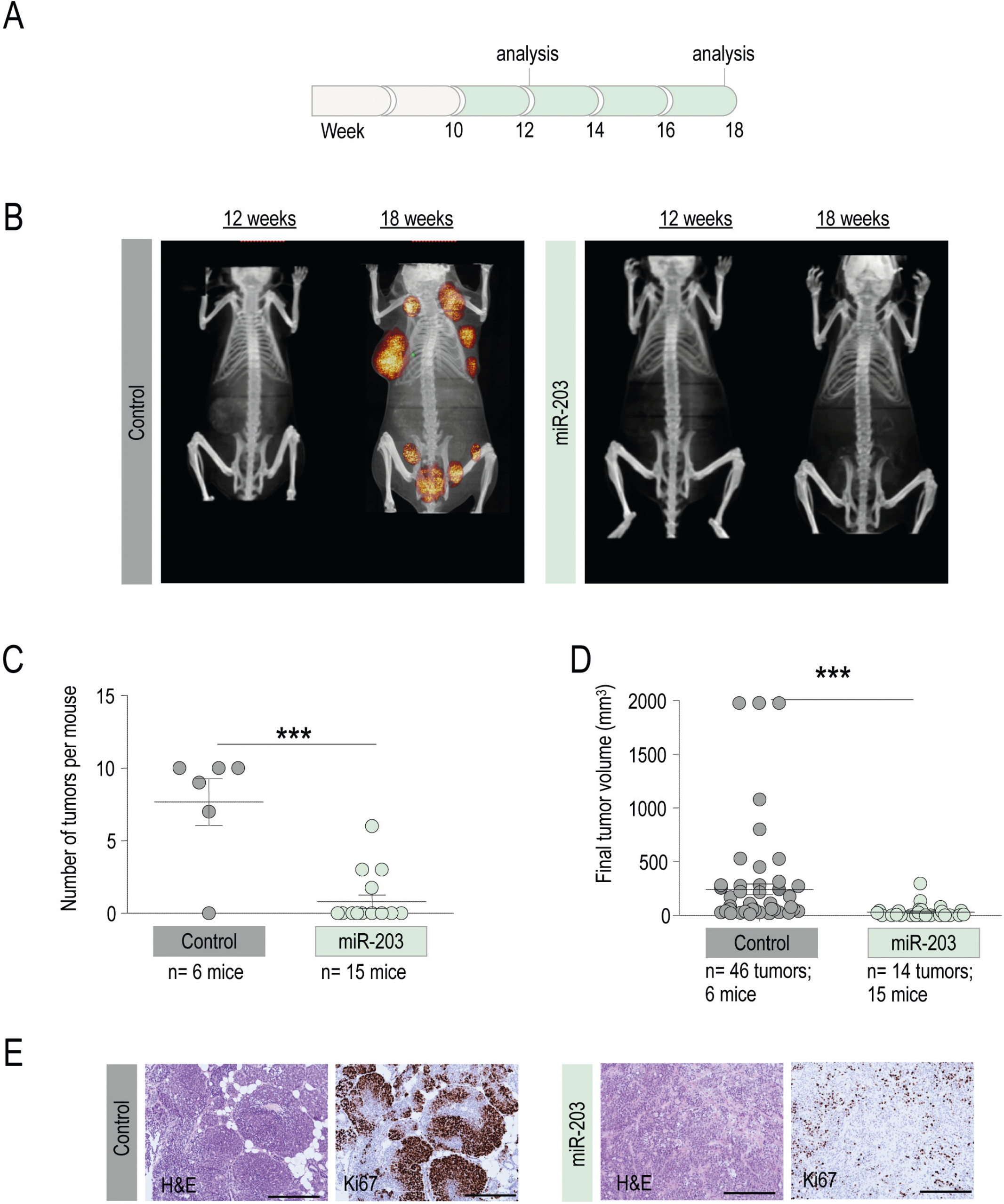
*In vivo* effects of miR-203 treatment on PyMT mice, started at tumor onset and sustained to the human endpoint. A, Schematic of the Dox treatment (in green) schedule *in vivo*, on *miR-203 wild-type* or *miR-203 knock-in*; *PyMT* mice, from week 10 to the experimental endpoint. B, Representative micro-CT images of mice subjected to the Dox treatment (in the figures, “control” indicates miR-203 wild-type; “miR-203” indicates knock-in mice), at 12 weeks of age and at the endpoint (18 weeks of age). C, Number of tumors per mouse at the endpoint, in control and miR-203-treated mice. D, Final tumor volume of control and miR-203-treated mice. In (C, D), data are represented as mean ± s.d. (number of mice and total number of tumors per group are indicated in the figure). ***p<0.001 (Student’s t-test). E, Representative H&E and Ki-67 IHC staining of control and miR-203-treated tumors at the endpoint. Scale bar, 500 μm.

**Fig. 3.**
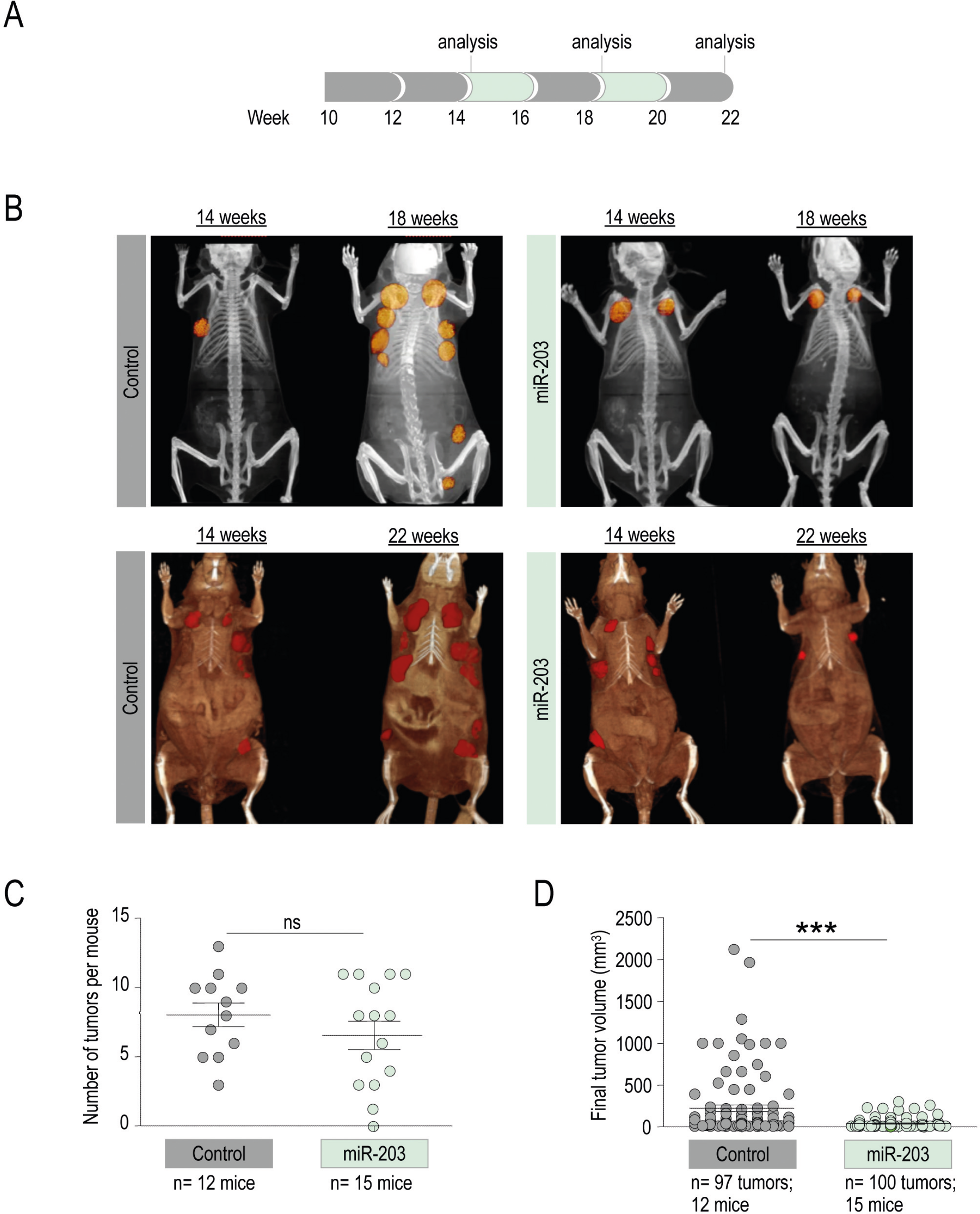
*In vivo* effects of miR-203 treatment on PyMT mice, started at tumor CT detection and administered every two weeks. A, Schematic of the Dox treatment (in green) schedule *in vivo*, on *miR-203 wild-type* or *miR-203 knock-in*; *PyMT* mice, starting when tumors are found by micro-CT imaging (around week 14) to the endpoint, on alternating weeks. B, Representative micro-CT images of mice subjected to the Dox treatment (in the figures, “control” indicates miR-203 wild-type; “miR-203” indicates knock-in mice) at tumor detection by micro-CT (14 weeks), four weeks later (18 weeks) and at the endpoint (22 weeks). C, Number of tumors per mouse at the endpoint, in control and miR-203-treated mice. D, Final tumor volume of control and miR-203-treated mice. In (C, D), data are represented as mean ± s.d. (number of mice and total number of tumors per group are indicated in the figure). ***p<0.001; *n.s*. not statistically different (Student’s t-test).

As depicted in Fig.1, we first treated *Tg (MMTV-PyMT); ColA1 (miR-203/miR-203); Rosa26 (rtTA/rtTA)* mice (for short, *PyMT; miR-203 wildtype* or *PyMT; miR-203 knock-in*) with Dox during two weeks from tumor onset (week 10, as detected by hematoxylin and eosin staining; Fig. 1A, Supplementary Fig. S1), followed by Dox withdrawal to the experimental human endpoint. The incidence of tumors per mice and the final tumor volume were significantly reduced in miR-203-treated compared to control mice (Fig.1B-1D). When the tumor samples were analyzed by immunohistochemistry at the end of the experiment, we found a down-regulation of the proliferation marker Ki-67 in those tumors briefly treated with miR-203 *in vivo*, suggesting a less aggressive phenotype (Fig.1E).

As a next step, we maintained the Dox treatment from tumor onset to the experimental human endpoint (Fig. 2A, Supplementary Fig. S1). The miR-203-mediated anti-tumor effects were stronger in this case, since miRNA treatment blocked tumor onset almost completely (Fig. 2B, 2C). Indeed, very few tumors were found in the miR-203-treated group, and their final volume was notably reduced when compared to the control counterparts (Fig. 2C, 2D). The rare and small miR-203-exposed tumors we were able to analyze exhibited again a markedly reduced proliferation rate (Fig. 2E).

Finally, we tested an *in vivo* schedule where the treatment started once the tumors were detectable by micro-CT (around week 14) and was intermittently applied to mice, every two weeks, to the experimental endpoint (Fig. 3A). Of interest, this treatment schedule was significantly effective in terms of tumor growth control but the average of tumors detected per mouse was almost identical to the one observed in the control group (Fig. 3B-3D), suggesting that, at this stage, tumor initiation capacity was recovered when the exposure to miR-203 was discontinuous. Since tumor initiation capacity falls -by definition- on dedifferentiated tumor cells, we wondered whether well-established markers for dedifferentiation in cancer (48) had been altered by miR-203 treatment. CD44 and NeuN expression levels were notably reduced in those tumors exposed to miR-203 *in vivo*, as well as the proliferation marker Ki67 (Fig. 4A), while H3K27me3, prolactin and progesterone receptor, the three of them considered markers of maturation and differentiation (48–51), were induced on miR-203-treated tumors respect to the control counterparts (Fig. 4B). As an additional key observation, none of the miR-203-treated mice developed lung metastasis in any of the three *in vivo* experiments performed, compared to 31% incidence of lung metastasis in the control groups (Fig. 4C).

**Fig. 4.**
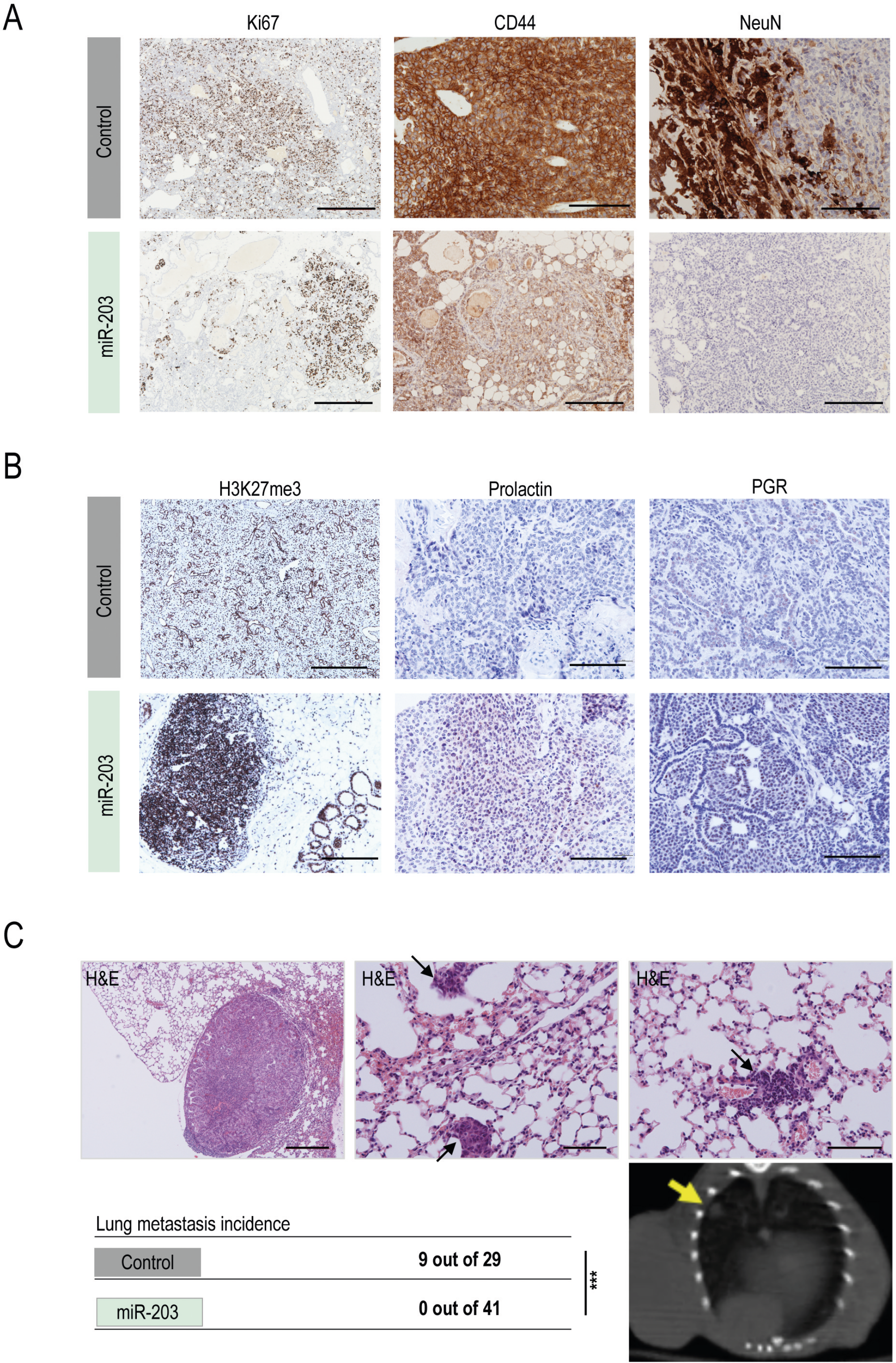
miR-203 exposure *in vivo* on PyMT mice alters the expression of stem-like and differentiation markers in mammary tumors, and fully prevents lung metastasis. A, Representative images of IHC staining for Ki67 (to test proliferation), CD44 and NeuN (as stem-like cell markers) in control and miR-203-treated tumors, at the experimental endpoint and after exposure to Dox on alternating weeks from tumor detection by micro-CT, as indicated in Fig. 3A. B, Representative images of IHC staining for H3K27me3, prolactin and progesterone receptor (PGR), to test evidences of differentiation on control and miR-203-treated tumors as in A. C, Illustrative H&E staining of lung macro- and micro-metastasis, found in several control mice at the experimental human endpoint. Representative examples are shown, from the 9 metastasis cases identified throughout the three *in vivo* experiments (depicted in Fig. 1, 2, 3). As shown in the table, the overall incidence of metastasis was 31,03% in control mice *versus* 0% in miR-203-treated mice. ***p<0.001 (Student’s t-test). The bottom right panel shows a representative micro-CT image, pointing to one evident macro-metastasis (yellow arrow) found in a control mouse. Scale bar, 500 μm.

Altogether, these observations demonstrate a beneficial role of miR-203 treatment not only to block proliferation and induce exhaustion of tumor-growth capacity, but also to ameliorate metastasis incidence. With these data, it is tempting to speculate that miR-203 treatment has an impact on the cell renewal and plasticity of cancer cells.

### Brief exposure to miR-203 induces morphological and molecular changes suggestive of epithelial differentiation on PyMT mammary tumor-derived organoids

Recently, the culturing of mammary organoids in 3D artificial extracellular matrix (ECM) hydrogels has gained popularity over 2D cell culture approaches, especially for studying mammalian development, disease and stem cell behavior (52). Previous studies demonstrated that organoids developed from breast tumors closely resemble the gene expression signature and heterogeneity of the tumor of origin, and even mammary branching morphogenesis is recapitulated in an organoid system by retaining its epithelial spatial organization (53–55). Therefore, we decided to create an organoid platform *in vitro*, which helped us to interpret our observations *in vivo* and to characterize the antitumor effect of miR-203 with special focus on cancer differentiation. To generate the organoid cultures, we followed the same schedule depicted in Fig. 3, including this time a second control group: apart from *PyMT; miR-203 wildtype* treated *in vivo* with Dox, we included a group of *PyMT; miR-203 knock-in* mice treated *in vivo* with vehicle. The outcome of both control groups was undistinguishable, corroborating that (i) Dox has no effect *per se* and (ii) the miR-203 inducible system is not leaky (41).

Interestingly, the morphology of control tumor-derived organoids was remarkably different to the one observed on miR-203-treated tumor-derived organoids. As depicted in Fig. 5A, the structure of control tumor-derived organoids was compacted, disorganized, dense and grape-shaped. However, the miR-203-treated tumor-derived organoids were predominantly cystic and structured, suggestive of a luminal epithelium (54). Moreover, the immune-histology analysis revealed lower proliferative rates (Ki-67 staining) in miR-203-treated tumor-derived organoids when compared to their control counterparts (Fig. 5A). When the control organoids (never exposed to Dox *in vivo*) were exposed to miR-203 *in vitro* for a short period of time (5 days, followed by miR-203 withdrawal for two more weeks), their morphology systematically changed in a gradual manner turning into hollow-cysts (Fig. 5B, Supplementary Fig. S2A), showing again that miR-203 treatment boosts the cyst-forming ability of mammary epithelial cells. Of interest, such capacity has been attributed to ALDH-positive progenitors (56). Accordingly, the expression levels of ALDH1/2 was notably diminished when the organoids were briefly exposed to miR-203 *in vitro* (Fig. 5C), suggesting the terminal differentiation of such progenitors. Ki-67 levels were also decreased in those organoids exposed to miR-203 *in vitro*, as happened to the organoids derived from miR-203-exposed tumors *in vivo* (Supplementary Fig. S2B). The cystic organoids exposed to miR-203 eventually collapsed (as denoted in the bright-field images of Fig. 5B and Supplementary Fig. 2A), while the control organoids were easily maintained *in vitro* for several passages during months.

**Fig. 5.**
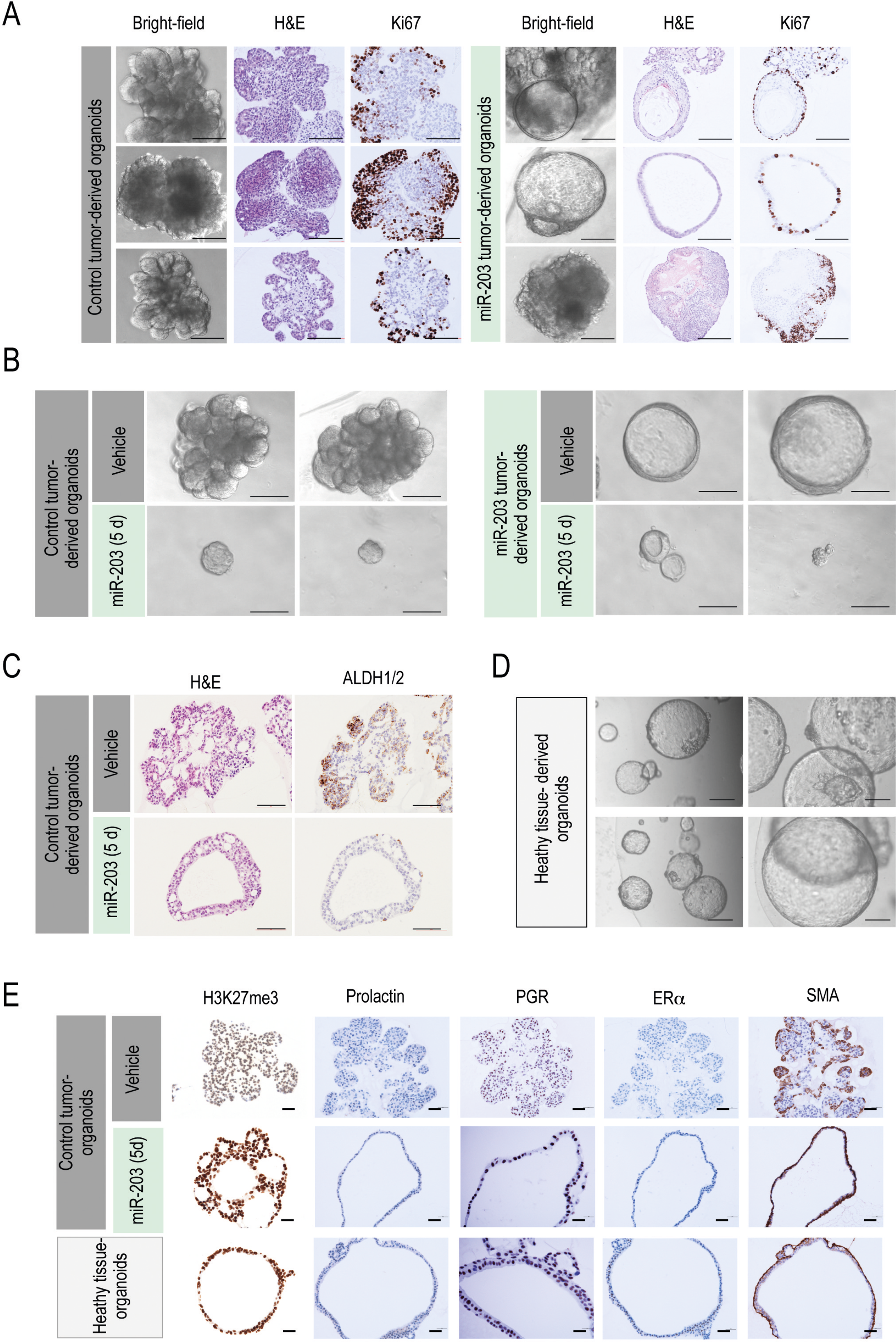
miR-203 transitory exposure promotes a morphological and molecular switch to epithelial differentiation on PyMT mammary tumor-derived organoids. A, Representative bright-field images and the corresponding H&E and Ki67 IHC staining of tumor-derived organoids (tumors from *miR-203 wild-type* or *miR-203 knock-in*; *PyMT* mice treated *in vivo* with Dox). B, Representative bright-field images of tumor-derived organoids (tumors from *miR-203 knock-in*; *PyMT* mice treated *in vivo* either with vehicle or Dox), exposed *in vitro* to vehicle or miR-203 (Dox) during 5 days and followed by miR-203 withdrawal for 2 more weeks (indicated as “miR-203 5d” in the figure). C, Representative images of ALDH1/2 IHC staining of control tumor-derived organoids, exposed to vehicle or miR-203 *in vitro* during 5 days and followed by miR-203 withdrawal, as in B. D, Representative bright-field images of healthy mammary gland-derived organoids (from *miR-203 wild-type*; *PyMT wild-type* mice). E, Illustrative IHC images of staining for H3K27me3, prolactin, progesterone receptor (PGR), estrogen receptor alpha (ERa) and smooth muscle actin (SMA) in control tumor-derived organoids (upper panels), control tumor-derived organoids treated *in vitro* with miR-203 during 5 days (middle panels) and healthy mammary gland-derived organoids (bottom panels). In A-D: scale bar 100 μm. In E: scale bar, 500 μm.

Intriguingly, healthy mammary gland tissue-derived organoids exhibited a similar phenotype to the one observed in the miR-203-treated organoids: cystic, organized and morphologically luminal-like (Fig. 5D). The induced expression of the epigenetic marker H3K27me3, associated to differentiation (49, 57), was prominent in miR-203-treated tumor organoids and healthy mammary gland-derived organoids, when compared to the control tumor counterparts (Fig. 5E). We further analyzed the expression of other markers linked to differentiation, such as prolactin, progesterone receptor, estrogen receptor alpha, and smooth muscle actin (49), and both the healthy tissue-derived organoids and the miR-203-treated tumor organoids exhibited comparable staining for all the molecular markers tested (Fig. 5E).

Since miR-203 induced morphological and molecular changes in the tumor-derived organoids suggestive of cancer cell differentiation, we next examined whether these changes were similar to the ones triggered by other well-known differentiation stimuli. Thus, we tested in our culture a defined epithelial differentiation media (detailed in methods section) and FGF2 treatment, described to induce branching morphogenesis (56, 58, 59). The healthy tissue-derived organoids mostly presented a cystic morphology in any condition tested, with the exception of FGF2 treatment, which always induced the mammary trees typical of branching morphogenesis (Supplementary Fig. S2C). On the contrary, the tumor-derived organoids were more susceptible to treatment-induced changes: while mostly condensed and grape-shaped upon basic expansion media, the tumor-derived organoids shifted to a predominant cystic morphology, induced by epithelial differentiation media and particularly by miR-203 expression, suggesting that any of those treatments were boosting the cyst-forming ability of mammary epithelial cells. Interestingly, and as occurred with the healthy tissue-derived organoids, those tumor organoids cultured in the presence of FGF2 exhibited a prominent branching morphology, either treated or not with miR-203 (Supplementary Fig. S2C).

Altogether, these observations suggest that a short exposure to miR-203 favors mammary epithelial differentiation on tumor organoids, which also correlates to a direct detrimental effect of this microRNA on the propagation and expansion of the organoid culture.

### Brief exposure to miR-203 induces a basal-to-luminal switch on mouse PyMT mammary tumor-derived organoids

In combination with other markers, cytokeratins (CK) have been used for a long time to determine the origin and grade of breast cancers. As represented in the schematic of Fig. 6A, cytokeratins 5, 14, and 17 are mostly associated to basal (and therefore poorly differentiated) tumors and poor patient prognosis, while cytokeratins 8 and 18 depict a luminal origin (and therefore highly differentiated status) and denote good patient prognosis (48, 57, 60–63). Following those well-established histopathology correlations, we tested by immunohistochemistry CK5, CK14 and CK8/18 expression levels in control tumors *versus* miR-203-treated tumors. Importantly, CK5 and CK14 staining were markedly reduced, while CK8/18 expression levels were induced in miR-203-treated tumors when compared to the control ones (Fig. 6B) suggesting a basal-to-luminal switch. Accordingly, tumor-derived organoids exhibited expression levels for CK5, CK14 and CK8/18 comparable to their corresponding tumors of origin, while miR-203 short exposure *in vitro* reduced the expression levels of CK5 and CK14 and induced the expression of CK8/18 (Fig. 6C). Again, the staining in miR-203-exposed organoids was comparable to that in healthy tissue-derived organoids for all the CK markers tested (Fig. 6C).

**Fig. 6.**
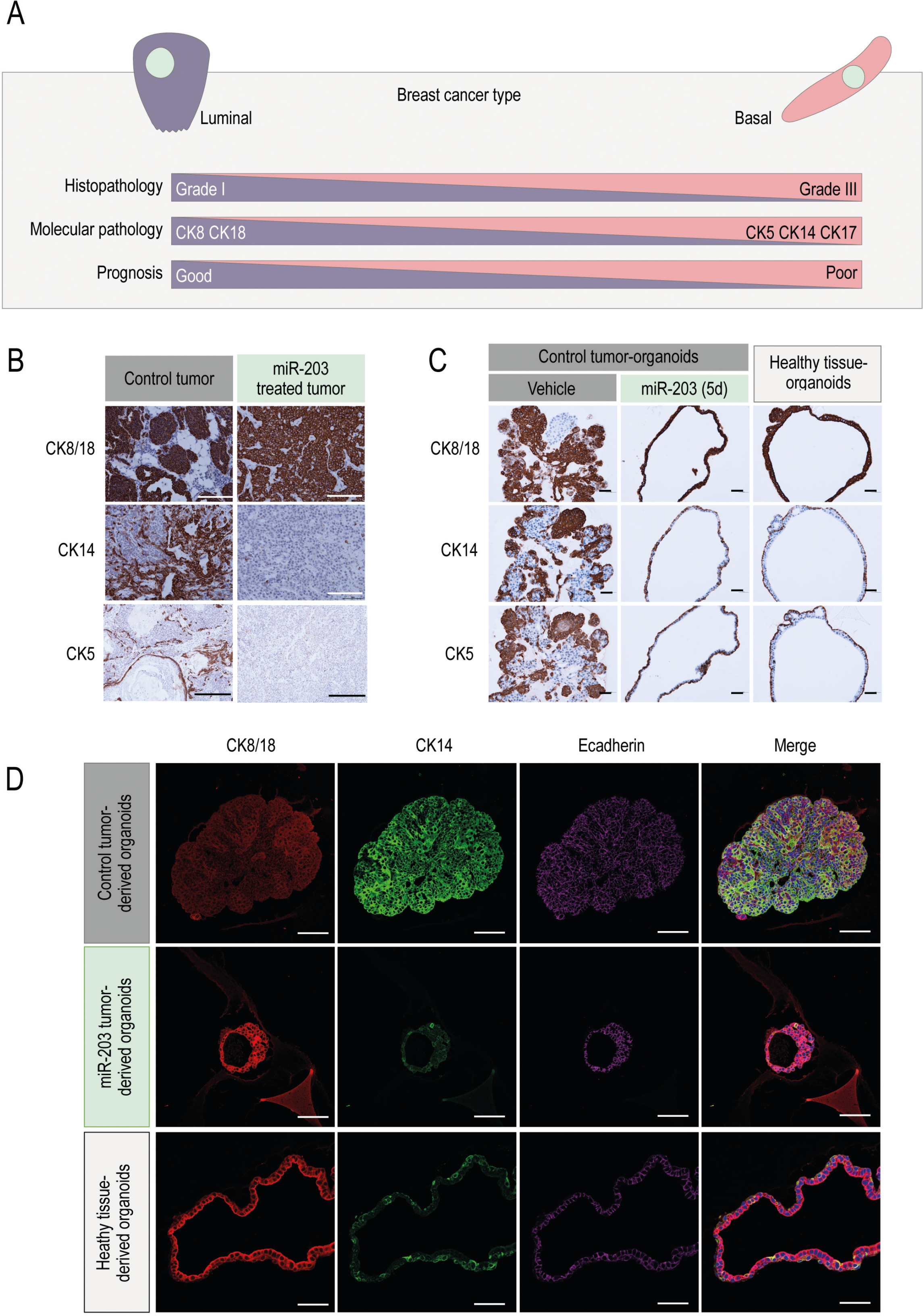
miR-203 transitory exposure induces a basal-to-luminal shift on mouse mammary tumor-derived organoids. A, Schematic showing the correlation between cytokeratins expression, histopathological tumor grade, prognosis and breast cancer type. B, Illustrative detection of CK8/18, CK14 and CK5 by IHC in control tumors and miR-203-treated tumors at the experimental endpoint. The doxycycline schedule followed for this set of experiments is also the one indicated in Fig. 3A. C, Illustrative IHC images of staining for CK8/18, CK14 and CK5 in control tumor-derived organoids (left panels), control tumor-derived organoids treated *in vitro* with miR-203 during 5 days (middle panels) and healthy mammary gland-derived organoids (right panels). D, Detection of CK8/18 (red), CK14 (green) and E-cadherin (purple) by immunofluorescence in tumor-derived organoids, extracted from control tumors (upper panels), miR-203-treated tumors (middle panels) or healthy mammary gland tissue (bottom panels) samples. In B-C: scale bar, 500 μm; in D: scale bar, 100 μm.

We performed additional immunofluorescence experiments on miR-203-exposed tumor-derived organoids and the corresponding controls, to corroborate those observations. The upper images in Fig. 6D show a representative example of control tumor-derived organoids, exhibiting high levels of CK14 and low levels of CK8/18, while miR-203-treated tumor-derived organoids (middle images) shifted the cytokeratins expression profile, being the CK8/18 the most predominant and CK14 becoming much less represented, almost absent. Again, the healthy tissue-derived organoids (lower images) exhibited a similar phenotype to the one observed in the miR-203-treated tumor-derived organoids.

To further understand the mechanistic insights of the differentiation-based antitumor effects evoked by miR-203, we performed RNA sequencing of organoid samples, derived from healthy or tumor tissue, and exposed *in vitro* to miR-203 (during 5 days, followed by 14 days of miR-203 withdrawal) or epithelial differentiation media (Supplementary Fig. S3). Principal Component Analysis of those samples revealed a prominent effect of miR-203 treatment only on tumor-derived organoids. To some extent, the transcriptomic profile induced by miR-203 on the tumor organoids appeared to be parallel to that induced by the differentiation media, suggesting a partial similarity between both treatments (Supplementary Fig. S3A). On the other hand, the differentiation media notably altered -while miR-203 treatment did not significantly modify- the transcriptomic profile of healthy tissue samples (Supplementary Fig. S3A). When a specific signature for “Mammary Gland Development” was considered, we identified a divergence between non-tumor and tumor organoids, as expected. Of interest, miR-203 treatment partially reverted such differences only in tumor organoids, for genes involved in this particular signature and also in the sub-cluster “Mammary Gland Epithelial Differentiation” (Supplementary Fig. S3B, S3C). Enrichment plots for the “Mammary Gland Stem Cell (MaSC)” and “Mature Luminal Cell” signatures revealed a significant correlation between miR-203 treatment and the induction of genes characteristic of mature luminal cells, while no correlation was observed for MaSC genes (Supplementary Fig. S3D). When we interrogated the gene expression profile of the Epithelial-to-Mesenchymal Transition (EMT), the miR-203 exposed samples showed a poor correlation, in contrast to those incubated in differentiation media (Supplementary Fig. S3D). Besides, the bulk RNA sequencing performed here supported our former observations, such as major alterations provoked by miR-203 brief exposure in the mRNA expression levels of cytokeratins [i.e. Krt14 (fold decrease: −2,3), Krt5 (fold decrease: −1,6), Krt18 (fold increase: 0,3)]; progenitor markers [i.e. Pygo2 (fold decrease: −0,3); Neat1 (fold decrease: −0,2); Per2 (fold decrease: −1,1); Cd44 (fold decrease: −0,7); Sox9 (fold decrease: −0,5); Klf4 (fold decrease: −1,6); Axin2 (fold decrease: −0,2); p63 (fold decrease: −0,2); Foxm1 (fold decrease: −0,7); Foxq1 (fold decrease: −0,5); HOTAIR (fold decrease: −0,8)]; EMT markers [i.e. N-cadherin (fold decrease: −0,3); Snai1 (fold decrease: −0,6); Snai2 (fold decrease: −0,8); Zeb2 (fold decrease: −0,3); Smad3 (fold decrease: −1,5)]; differentiation markers [Prolactin receptor (fold increase: 0,7); Gata3 (fold increase: 0,8); Atf3 (fold increase: 0,9); Myb (fold increase: 0,6)] and interestingly, in the mRNA expression levels of key cell cycle regulators, such as Cdk1 (fold decrease: −2,5), among others. Of interest, gene signatures for “Basal Cells” (as defined by two different data bases) were significantly down-regulated by miR-203 treatment (Supplementary Fig. S4A) as well as gene signatures for “Organ and Cell Development”, “Cell Migration and Motility”, “Cell Metabolism” and “Cell Cycle” (Supplementary Fig. S4B and Supplementary Table 1).

Altogether, these data on PyMT breast cancer organoids demonstrate that a brief exposure to miR-203, either *in vitro* or *in vivo*, induces a shift from a basal tumor phenotype to a more differentiated luminal epithelial status.

### Brief exposure to miR-203 induces a basal-to-luminal shift and reduces collective migration on patient-derived breast tumor organoids

To explore the therapeutic potential of miR-203 in humans, we evaluated its effects on breast cancer patient-derived organoids. Thus, more than 10 independent patient-derived 3D cultures were efficiently generated from cylinders of core-needle biopsies (to BIRAD 4C-5 patients). Fig. 7A shows a schematic of the procedures followed with patient-derived organoids and the temporal line of the experimental settings. As depicted, after 7 days of culture establishment and organoid amplification, patient-derived 3D cultures were transiently transfected with synthetic miR-203 mimics, followed by miR-203 withdrawal for three additional weeks, when the analysis was performed. We observed the 3D cultures under the bright-field microscope along the experiments to evaluate every potential morphological alteration induced by the short exposure to miR-203. Soon after organoid establishment, we systematically observed the formation of cell spire protrusions only in control organoids (Fig. 7B). Elongated cells emerged from the organoid edges and, whenever made contact with a solid surface (i.e. the plastic or glass well bottom), they attached to it and gradually occupied the surrounding area forming a bi-dimensional layer below and beyond the tridimensional organoids (Fig. 7B). It is reasonable to speculate that these cells undergo collective migration. It has been described that tumor cells may experience a partial EMT with their cell-cell connections remaining intact, and thereby migrate as a cohesive group (64). The leader cells use similar mechanisms as migrating single cells to polarize, protrude, invade and adhere to stromal matrix and they are generally more organized and efficient in direct invasion than the individual cells (64–66). Molecularly, we detected a reproducible pattern of front-rear polarity for the expression of cytokeratins and the EMT marker vimentin (Fig. 7C): in control organoids, CK14 and vimentin appeared highly expressed within the cells conforming the external organoid layer and those attaching to the plate surface, while CK8/18 was almost undetectable. Of interest, a short and transient exposure to miR-203 blocked the cell projections and migration from the organoids (images in Fig. 7C, 7D and quantification in Fig. 7D), reduced the expression of CK14 and vimentin to almost undetectable levels and in turn, stimulated the expression of CK8/18 (Fig. 7C). Of importance, not only collective migration was dropped by the exposure to miR-203 but also the total number of organoids, their complexity and their size were notably reduced, while the proportion of luminal-like organoids in the culture was significantly augmented (Fig. 7D).

**Fig. 7.**
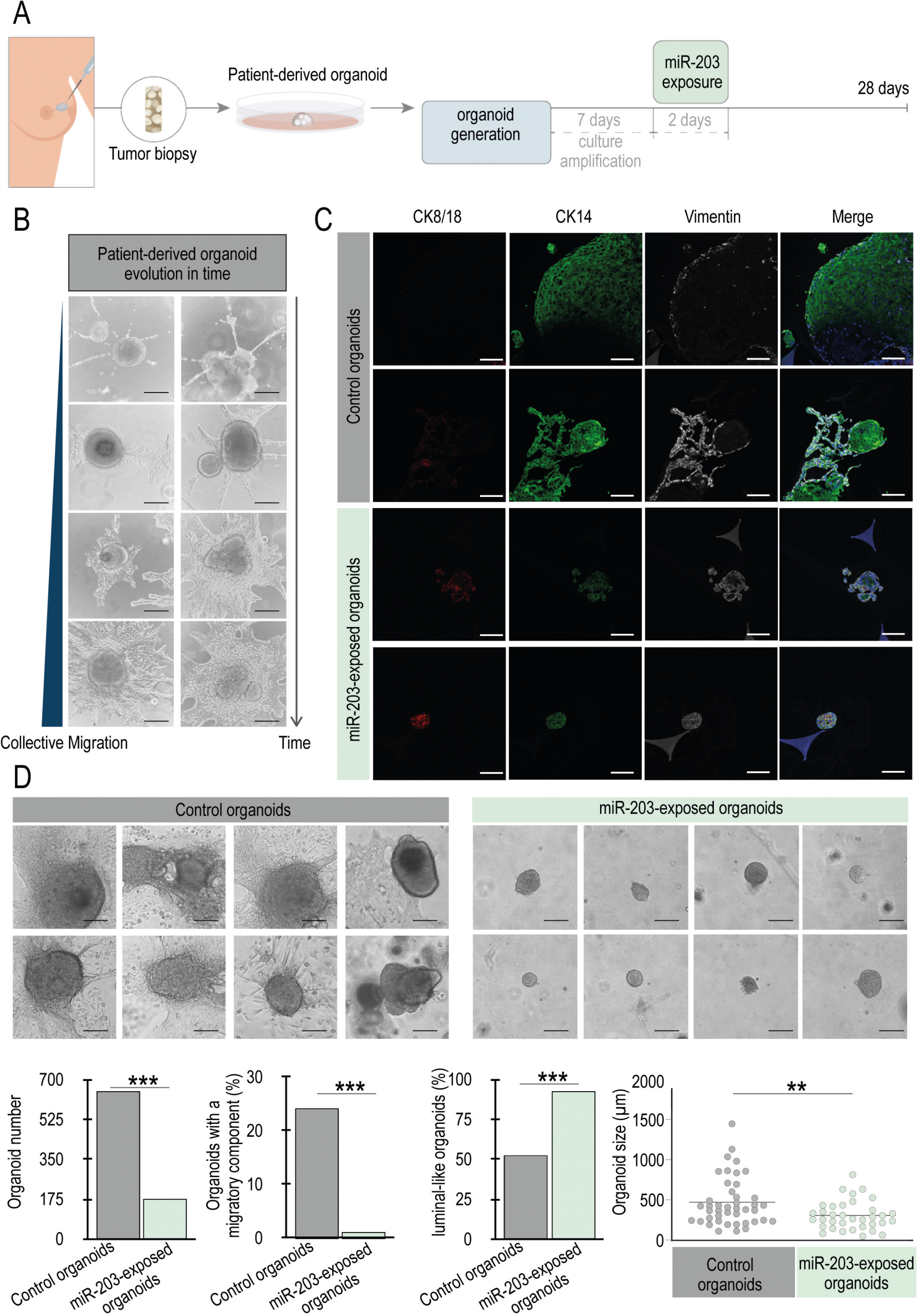
miR-203 transitory exposure induces a basal-to-luminal shift and reduces collective migration on patient-derived breast tumor organoids. A, Schematic showing the experimental procedures followed for patient-derived tumor processing, organoid culture establishment and miR-203 mimics transient transfection. B, Representative bright-field images showing the progressive collective cell migration projected from the 3D patient-derived organoids along time. C, Detection of CK8/18 (red), CK14 (green) and vimentin (white) by immunofluorescence in patient tumor-derived organoids, transiently exposed or not to miR-203 mimics *in vitro*. D, Representative bright-field images of patient-derived organoids, control *versus* miR-203 briefly exposed, denoting the morphological differences in complexity, size and migration upon miR-203 treatment. Lower panels: quantification of the total number of organoids, percentage of organoids exhibiting collective migration, percentage of organoids with luminal-like morphology and organoid size, of control *versus* miR-203 briefly exposed patient-derived organoids. **p<0.01; ***p<0.001 (Student’s t-test), n=3 biopsy-derived organoid cultures. In B-D: Scale bar, 100 μm.

Altogether, the data presented here corroborate, in patient-derived samples, our previous observation in murine samples and thus reinforce the potential of miR-203 as a cancer differentiation driver.

## Discussion

Cancer has been broadly interpreted as a caricature of normal tissue development. Cellular programs regulating tissue plasticity, self-renewal and expansion are exquisitely orchestrated under physiological conditions. However, aberrant tumor mechanisms unbalance this coordinated cell plasticity and give rise to immature or dedifferentiated tumor cells. Indeed, it is now widely accepted that, for tumor initiation, adult cells experience reprogramming to a progenitor-like fate (67). Thus, tumor dedifferentiation supports cancer progression, relapse and metastasis. Traditional chemo and radiotherapy generally involves the elimination of proliferating tumor cells. Instead, the differentiation therapies offer the possibility of coaxing cancer cells into becoming normal cells, reactivating the endogenous differentiation programs to resume maturation. Although promising, the cancer differentiation approaches are still evolving and require novel methodologies to reach efficient therapies. Trusting this general believe, we decided to examine the antitumor effects of miR-203 from a cancer differentiation perspective. This microRNA has been recently described by our group to fine-tune the critical balance between reprogramming, stemness and differentiation programs: miR-203 blocks somatic-to-pluripotency reprogramming (40) while potentiates differentiation of stem cells to a mature and terminal state (41). We wondered whether those effects would be applied to cancer differentiation and therefore would point to miR-203 as a promising tool for differentiation-based antitumor therapy.

*In vivo*, we tested different treatment schedules, trying to understand the consequences of treatment intermittency. Interestingly, only when mice were exposed to miR-203 from tumor onset to the end of the experiment, tumor initiation and growth were completely prevented. When miR-203 treatment was intermittent, we noticed a significant control on tumor growth and a considerable delay in tumor initiation, while we did not completely avoid the latter. In this line, recent advances on stem cell biology have demonstrated that stem cell plasticity represents one of the major therapeutic challenges for differentiation therapies. Several studies have provided evidence that both CSCs and non-CSCs are plastic and capable of undergoing phenotypic transitions in response to appropriate stimuli. This notion was for instance exemplified by a study in which cell populations displaying stem cell–, basal-, or luminal-like phenotypes were isolated from breast cancer cell lines (68). *In vitro*, all three subpopulations were able to generate cells of the other two phenotypes. This phenotypic inter-conversion was stochastic and not determined by the cell phenotype of origin. Thus, it is accepted now that CSC and non-CSC states are not hardwired: considering that plasticity may be in tumor cells as extensive as it is in healthy tissues, CSCs would be always recreated. This fact could explain the successful outcome when miR-203 treatment was uninterrupted (Fig. 2), theoretically capturing any newly formed CSC. However, the intermittency in miR-203 exposure would tentatively allow the undifferentiated population to restore, favoring then the tumor initiation process. Thus, we could speculate that miR-203 exposure was able to maintain the stem-like capacity delimited and therefore, tumor growth under control in any regimen tested. One of the key results *in vivo*, denoting the strong effect of miR-203 on tumor differentiation, was the lack of metastasis (suggestive of detrimental migratory and invasive capacity) detected on those mice exposed to miR-203 in any of the treatment schedules tested, while their control counterparts experience a metastasis incidence of 31%. Of interest, *in vitro* cell cultures derived from patient biopsies exhibited a polarized and migratory cell population, that was completely abolished by miR-203 treatment. Together, those two interesting observations prompt us to speculate that miR-203 impact on cancer cell differentiation plays a role in invasion and metastasis.

Inspired by three-dimensional cell culture systems of the mammary epithelium, Hans Clevers and collaborators established culture protocols that allow the generation and long-term expansion of three-dimensional epithelial organoids (69, 70). Those adult stem cell-based cultures lack mesenchymal cells and generally require the following niche factors: the mitogen epidermal growth factor (EGF), the Wnt-agonist R-spondin, the transforming growth factor beta (TGF-b) inhibitor Noggin, and the extracellular matrix surrogate basement membrane extract (BME, or Matrigel). Depending on species and organ, additional common additives are, among others, Wnt-3A and fibroblast growth factor (FGF) 10, the activin receptor-like kinase (ALK) inhibitor A83-01, the p38 mitogen-activated protein kinase inhibitor SB202190, and the vitamin nicotinamide (54, 70). Thus, following Clevers’ protocols, we efficiently produced organoids from the mouse tumors or mammary glands, recapitulating *in vitro* the tridimensional architecture and the molecular features of the source tissue.

Our first striking observation was the remarkable differences on organoid morphology upon miR-203 treatment: the structure of control tumor-derived organoids was notably compacted, disorganized, dense and in most cases, grape-shaped, while the miR-203-treated tumor-derived organoids were predominantly cystic, suggesting an epithelial luminal origin. Of interest, the cystic morphology was reproduced any time the tumor-derived organoids were yielded to transient miR-203 treatment *in vitro*. Those cystic organoids resulting from miR-203 exposure, either *in vivo* or *in vitro*, collapsed after a few passages, contrary to the control organoids never treated with the microRNA, which were long-term maintained in culture, as published before (54, 69, 70). The short lifespan of organoids exposed to miR-203 clearly pointed to an exhaustion of the self-renewal capacity of the culture and implied a direct detrimental effect by this miRNA on the propagation and expansion of the organoids. Accordingly, Ki-67, CK5, CK14 and ALDH1/2 levels were diminished in those miR-203-exposed organoids, while markers such as CK8/18 were induced, denoting a shift from dedifferentiated-basal to a more differentiated luminal-like phenotype. Even more, the healthy mammary gland tissue-derived organoids exhibited a very similar phenotype to the one observed in the miR-203-treated tumor organoids. Several markers associated to differentiation were found to be more expressed in healthy tissue-derived as well as in miR-203-exposed tumor organoids. Deepening into the differentiation concept, we tested in our organoid platform some previously defined differentiation conditions (such as FGF2-mediated induction of mammary branching or a culture medium described for mammary epithelial cell differentiation). The tumor-derived organoids were mostly condensed and grape-shaped upon basic expansion media, and shifted to a predominantly hollow-cysts morphology when exposed to the epithelial differentiation media or (even more dramatically) to miR-203, suggesting that any of those treatments were boosting the cyst-forming ability of mammary epithelial cells. As expected, FGF2 treatment always induced the mammary trees typical of branching morphogenesis (56) either on non-tumor or tumor organoids, exposed or not to miR-203. The intriguing fact that, upon FGF2 treatment, miR-203 does not induce epithelial cystic formations but instead favors the branching constructions, was in concordance with our previous works (40, 41) and suggests that, submitted to a strong differentiation scenario, miR-203 acts always as a differentiation enhancer, never as a reprogramming inducer.

Importantly, this molecular and phenotypical analysis of the organoids have uncovered the miR-203 effects on all the three axes governing cancer cell differentiation in breast tumors: i.e. dedifferentiated-to-mature; mesenchymal-to-epithelial; basal-to-luminal. The three are certainly inter-connected although not responding to identical molecular pathways. The fact that a brief exposure of cancer cells to miR-203 alters the expression of markers widely associated to the three axis reinforces our message and points to miR-203 as an encouraging tool for cancer cell differentiation.

When the transcriptomic profiles of those organoids were tested by RNA sequencing we noticed a remarkable impact of miR-203 treatment on the differentiation of tumor-derived organoids, while the healthy tissue-derived organoids showed very little alterations when exposed to miR-203. These data were in concordance with the phenotypically observed modifications incited by miR-203: whereas miR-203 exposure completely shifted the shape of tumor organoids, no notable changes were induced on non-tumor organoids. This suggests a fascinating differential impact of this microRNA on tumor and non-tumor tissues that deserves to be further explored through single cell sequencing, at the transcriptomic, genomic and epigenetic levels.

Altogether, the data presented here confirm the antitumor effects mediated by miR-203, particularly denoting now its influence on cancer differentiation, both in murine and patient-derived samples. This work undoubtedly opens new perspectives on the potential therapeutic applications of miR-203 in cancer.

## Methods

### Animal models and procedures

Animal experimentation was performed according to protocols approved by the CNIO-ISCIII and the UCM Ethics Committee for Research and Animal Welfare (CEIyBA) and Madrid Regional Government, according to European official regulations. The miR-203 inducible model was generated by cloning a 482-bp genomic mmu-mir203 sequence into the pBS31 vector for recombination into the ColA1 locus in embryonic stem cells. The resulting knock in allele [ColA1(miR-203)] was combined with a Rosa26-M2rtTA allele [Rosa26(rtTA)] for doxycycline-dependent induction as described previously (41). PyMT [FVB/N-Tg(MMTV-PyVT)^634Mul/J^] mice were kindly provided by Miguel Quintela (CNIO, Spain). Mice were then crossed to obtain the Tg (MMTV-PyMT); Rosa26(rtTA); ColA1(miR-203) strain, which has been used throughout this work. To induce miR-203 expression *in vivo*, Doxycycline (Dox) was orally administered to mice in diet (Dox-delayed release pellets, from Jackson laboratories) following the different schedules indicated in Figures 1–3. As control, Dox treatment was applied to Tg (MMTV-PyMT); miR-203 (+/+) mice, which also served as an internal checkup of the Dox treatment itself.

Primers used for genotyping the PyMT transgene were 5′-GGAAGCAAGTACTTCACAAGG-3′ and 3′-GGAAAGTCACTAGGAGCAGGG-5′. Polymerase chain reaction conditions were as follows: 95 °C for 15 min; 94 °C for 30 s; 30 cycles at 59 °C for 45 s; 72 °C for 1 min; 72 °C for 10 min; then soaking at 4 °C. PCR products are 336 bp (base pair) for wt allele, 438 bp for lox allele, 470 bp for cre allele, and 557 bp for PyMT allele. All these animals were maintained in a mixed C57BL6/J x 129 x CD1 genetic background and were housed at the serum pathogen free (SPF) barrier area of the CNIO. Mice were treated in accordance with the Spanish Laws and the Guidelines for Human Endpoints for animals used in Biomedical Research. Mice were observed on a daily basis and sacrificed when they showed signs of morbidity or overt tumors.

### Micro computed tomography (micro-CT)

For micro-CT, mice were anesthetized with a continuous flow of 1% to 3% isoflurane/oxygen mixture (2 L/min). Acquisitions were performed using a micro-CT scanner Argus-Vista (SEDECAL, Madrid, Spain) including the whole body in 2-bed position. Tomographic images were reconstructed using a 3D-FBP (filtered back projection) algorithm that produced 55 slices measuring 55 × 55 pixels each. The isotropic resolution of this instrument was 45 μm. The micro-CT image acquisition consisted of 400 projections collected in one full rotation of the gantry in approximately 10 min per bed position. The image acquisition was made without any contrast agent. The X-ray tube settings were 80 kV and 450 μA. For image analysis and quantification, 3D Slicer software was used. Tumor volumes were measured once per week by micro-CT, to determine accurately the tridimensional tumor mass. The investigators were blinded during the entire *in vivo* experiment (micro-CT measurements were performed in all cases with no information about the genotype or treatment of every mouse tested). The potential effects of miR-203 on metastasis incidence in the lungs were also analyzed by micro-CT throughout the three *in vivo* experiments.

### Mammary gland-derived organoids culture

*Tg (MMTV-PyMT); miR-203 (+/+)* and *Tg (MMTV-PyMT); miR-203 (KI/KI)* mice (treated or not with Dox *in vivo*, as indicated in the text) were euthanized, and tumors were extracted. Two random pieces were snap frozen and stored at −80°C; two random pieces were fixed in formalin for histopathology and immunohistochemistry analysis and the remainder was processed for the isolation of viable cells. The remaining tissue was minced, washed with 10 mL AdDF+++ (Advanced DMEM/F12 containing 1x Glutamax, 10 mM HEPES, and antibiotics) and digested in 10 mL <u>BC organoid expansion medium</u>: 10% homemade R-Spondin 1 conditioned medium; 5 nM neuregulin 1 (Peprotech 100-03); 5 ng/mL FGF7 (Peprotech 100-19); 20 ng/mL FGF10 (Peprotech 100-26); 5 ng/mL EGF (Peprotech AF-100-15); 100 ng/mL Noggin (Peprotech 120-10C); 500nM A83-01 (Tocris 2939); 5 μM Y-27632 (Abmole); 500 nM SB202190 (Sigma S7067); 1X B27 supplement (Gibco 17504-44); 1,25 mM N-Acetylcysteine (Sigma A9165); 5mM nicotinamide (Sigma N0636;) 1X Glutamax (Invitrogen 12634-034); 10mM Hepes (Invitrogen 15630-056); 100U/mL Penicillin/Streptomycin (Invitrogen 15140-122); Primocin (Invitrogen Ant-pm-1) and Advanced DMEM/F12 (Invitrogen 12634-034), containing 1-2 mg/mL collagenase (Sigma, C9407). Digestion was performed on an orbital shaker at 37°C for 1-2 h. The digested tissue suspension was sequentially sheared using 10 mL and 5 mL plastic and flamed glass Pasteur pipettes. After every shearing step the suspension was strained over a 100 μm filter with retained tissue pieces entering a subsequent shearing step with ~ 10mL AdDF+++. 2% FCS were added to the strained suspension before centrifugation at 400 rcf. The pellet was resuspended in 10mL AdDF+++ and centrifuged again at 400 rcf. In case of a visible red pellet, erythrocytes were lysed in 2 mL red blood cell lysis buffer (Roche, 11814389001) for 5 min at room temperature before the addition of 10mL AdDF+++ and centrifugation at 400 rcf. The pellet was resuspended in 10 mg/mL cold Cultrex growth factor reduced BME type 2 (Trevigen, 3533-010-02) and 40 μL drops of BME-cell suspension were allowed to solidify on pre-warmed 24-well suspension culture plates (Greiner, M9312) at 37°C for 20 min. Upon completed gelation, 400 μL of BC organoid expansion medium was added to each well and plates transferred to humidified 37°C / 5% CO_2_ incubators. Medium was changed every 4 days and organoids were passaged every week: organoids were resuspended in 2 mL cold AdDF+++ and mechanically sheared through flamed glass Pasteur pipettes. When necessary, very dense organoids were dissociated by resuspension in 2 mL TrypLE Express (Invitrogen, 12605036), incubation for 1-5 min at room temperature, and mechanical shearing through flamed glass Pasteur pipettes. Following the addition of 10 mL AdDF+++ and centrifugation at 300 rcf. or 400 rcf. respectively, organoid fragments were resuspended in cold BME and reseeded as above at ratios (1:1 to 1:6) allowing the formation of new organoids. Single cell suspensions were initially seeded at high density and reseeded at a lower density after ~ 1 week. In order to prevent misidentification and/or cross-contamination of BC organoids, we cultured every line physically separate. All organoid lines were frequently tested and resulted in all cases negative in the MycoAlert mycoplasma detection kit (Lonza, LT07-318). For epithelial differentiation, we used the media defined by Lonza (MEGM Mammary Epithelial Cell Growth Medium and Bullekit). Basically, this media has been optimized for the growth of mammary epithelial cells in a serum-free environment and includes BPE, hEGF, insulin, hydrocortisone and GA-1000 (Lonza CC-3150). FGF2 treatment (2 nM; Sigma) was used to induce mammary branching as published before (56). For inducing transient miR-203 over-expression, ColA1(miR-203/miR-203); Rosa26(rtTA/rtTA) organoid cultures were treated with Dox (1 μg/mL; Invitrogen) during 5 days. After that, Dox withdrawal was standardized for the cultures during following several passages (usually 2 weeks) unless other time points are indicated in the text. In this inducible system, we always test that insert expression is uniquely dependent on Dox, and becomes absolutely undetectable after Dox withdrawal. As a control of the treatment itself, Dox was also added and tested in wild-type organoids.

### Patient-derived organoids generation and culture

For this study, breast cancer patients (with BIRAD 4C-5-6) from Hospital “12 de Octubre” (Madrid, Spain) donated one cylinder of the first core-needle tumor biopsy, prior diagnosis. To guarantee the protection of patients enrolled in this study, we have strictly followed the hospital guidance, the local regulations, the “Declaration of Helsinki” and the Guidelines of good clinical practice from the “International Conference on Harmonization” ICH E6 (R2), effective from June 14, 2017. The technical protocols for patient-derived sample collection and processing and any additional material delivered to the patient (such as Patient Information Sheets or the Informed Consent Document), were carefully evaluated and approved by the corresponding Clinical Research Ethics Committee, in accordance with national legislation. Tumor samples were immediately processed in our laboratory for organoid culture generation, as described above. We were able to maintain patient-derived organoid cultures for three or four passages and the experiments were always performed at passage one. After 7 days of culture establishment and organoid amplification, patient-derived 3D cultures were transiently transfected with miR-203 mimics, followed by miR-203 withdrawal for three additional weeks. Hsa-miR-203 mimics were purchased from Sigma Aldrich (MISSION microRNA mimics) and transient transfection was performed using Lipofectamine 2000 (Sigma), following manufacturer’s instructions. Since then, cultures were carefully evaluated under the bright-field microscope for quantification of organoid number and size, complexity, formation of 2D projections, and finally, immunofluorescence was performed at the end of the experiment (three weeks after the miR-203 brief exposure).

### Immunofluorescence and immunohistochemistry

Organoids were fixed in 4% paraformaldehyde for at least 15 min, permeabilized using PBS 0.1% Triton X-100 for 15 min and blocked in BSA for 1 h at room temperature. Primary antibody incubation was performed overnight at 4°C in all cases, followed by secondary antibody incubation for 1 h at room temperature. Nuclear staining was included in the last PBS wash, using Hoescht or DAPI. Primary antibodies used in this study were against CK8/18 (rabbit monoclonal EP17/EP30, Dako, IR094), CK14 (rabbit polyclonal AF64, Covance, PRB-155P) and E-cadherin (mouse monoclonal 36, BD Bioscience, 610182) for mouse-derived samples and CK8/18 (rat monoclonal, DSHB, 531826), CK14 (rabbit monoclonal, Abcam, ab181595) and vimentin (mouse monoclonal RV202, BD Pharmigen, 550513) for patient-derived samples. Cells were examined under a Leica SP5 microscope equipped with white light laser and hybrid detection.

For immunohistochemistry, tissue samples were fixed in 10% neutral buffered formalin (4% formaldehyde in solution), paraffin-embedded and cut at 3 μm, mounted in superfrost®plus slides and dried overnight. Consecutive sections were stained with hematoxylin and eosin (H&E) or subjected to immunohistochemistry using automated immunostaining platforms (Ventana Discovery XT, Roche or Autostainer Plus Link 48). Antigen retrieval was first performed with high or low pH buffer depending on the primary antibody (CC1m, Roche or low pH antigen retrieval buffer, Dako), endogenous peroxidase was blocked (peroxide hydrogen at 3%) and slides were incubated with primary antibodies against Ki-67 (rabbit monoclonal D3B5, Cell Signalling Technology, 12202), CK5 (rabbit polyclonal AF 138, Covance, PRB-160P), SOX-10 (goat polyclonal N20, Santa Cruz Biotechnology, sc-17342), CD44 (rabbit polyclonal, Abcam, ab157107), H3K27me3 (rabbit monoclonal C36B11, Cell Signalling Technology, 9733), prolactin (rabbit polyclonal, Dako, A0569), progesterone receptor (rabbit monoclonal SP2, Thermo Scientific, RM-9102-R7), NeuN (mouse monoclonal A60, Millipore, MAB377), E-cadherin (mouse monoclonal 36, BD Bioscience, 610182), Aldh1/2 (mouse monoclonal H-8, Santa Cruz Biotechnology, sc-166362), CK8/18 (rabbit monoclonal EP17/EP30, Dako, IR094), CK14 (rabbit polyclonal AF64, Covance, PRB-155P), smooth muscle actin (mouse monoclonal 1A4, Dako, IR611), estrogen receptor alpha (rabbit polyclonal, Santa Cruz Biotechnology, sc-542).

Secondary antibodies were conjugated with horseradish peroxidase (OmniRabbit, Ventana, Roche) and the immunohistochemical reaction was developed using 3,30-diaminobenzidine tetrahydrochloride (DAB) as a chromogen (Chromomap DAB, Ventana, Roche or DAB solution, Dako) and nuclei were counterstained with Carazzi’s hematoxylin. Finally, the slides were dehydrated, cleared and mounted with a permanent mounting medium for microscopic evaluation. The images were acquired with a slide scanner (AxioScan Z1, Zeiss). Images were captured and quantified using the Zen Software (Zeiss).

### Analysis of mRNA levels, RNA sequencing

RNA/microRNA was extracted from organoids samples with Trizol (Invitrogen) or by using the miRVana isolation kit (Thermo Fisher), following the manufacturer’s recommendations and after the dissociation of matrigel/BME from the cultures by using the Cell Recovery Solution (Corning), following the manufacturer’s protocols. For reverse transcription of microRNAs, we used the Taqman small RNA assay (4366596), including the specific oligonucleotides for mmu-miR-203-5p and 3p (002580 and 000507), miR-16 and the housekeeping RNAs sno-202 or sno-142. Conditions for miRNA amplification were as follows: 30 minutes at 16°C; 30 minutes at 42°C and a final step of 5 minutes at 85°C. Quantitative real time PCR was then performed using the Taqman Universal PCR Master Mix (434437) following the manufacturer’s instructions in an ABI PRISM 7700 Thermocycler (Applied Biosystems).

For RNAseq, total RNA was extracted using the miRVana miRNA isolation kit (ThermoFisher), following the manufacturer’s recommendations. Between 0.8 and 1 μg of total RNA were extracted from organoids after dissociating the Matrigel/BME form the cultures (as indicated above). RIN (RNA integrity number) numbers were always in the range of 9 to 10 (Agilent 2100 Bioanalyzer). 250ng of total RNA samples were used. Average sample RNA Integrity Number was 9.1 (range 8.2-9.8) when assayed on an Agilent 2100 Bioanalyzer. Sequencing libraries were prepared with the “QuantSeq 3‘ mRNA-Seq Library Prep Kit (FWD) for Illumina” (Lexogen, Cat.No. 015) by following manufacturer instructions. This kit generates directional libraries stranded in the sense orientation, the read1 (the only read in single read format) has the sense orientation. Library generation is initiated by reverse transcription with oligodT priming, and a second strand synthesis is performed from random primers by a DNA polymerase. Primers from both steps contain Illumina-compatible sequences. Libraries were completed by PCR, applied to an Illumina flow cell for cluster generation and sequenced on an Illumina HiSeq 2500 with v4 Chemistry by following manufacturer’s protocols. Read adapters and polyA tails were removed with bbduk.sh (https://sourceforge.net/projects/bbmap/), following the Lexogen recommendations. Processed reads were analysed with the nextpresso pipeline (71), as follows: Sequencing quality was checked with FastQC v0.11.7 (http://www.bioinformatics.babraham.ac.uk/projects/fastqc/). Reads were aligned to the mouse reference genome (GRCm38) with TopHat-2.0.10 (72) using Bowtie 1.0.0 (73) and Samtools 0.1.19 (74) (library-type fr-secondstrand in TopHat), allowing two mismatches and twenty multihits. Read counts were obtained with HTSeq-count v0.6.1 (75) (stranded=yes), using the mouse gene annotation from GENCODE (gencode.vM20.GRCm38.Ensembl95). Differential expression was performed with DESeq2 (76), using a 0.05 FDR. GSEA Pre-ranked (77) was used to perform gene set enrichment analysis for several gene signatures on a pre-ranked gene list, setting 1000 gene set permutations. Only those gene sets with significant enrichment levels (FDR q-value < 0.25) were considered.

### Statistics

Samples (organoids or mice) were allocated to their experimental groups according to their pre-determined type and therefore there was no randomization. Investigators were blinded to the experimental groups in all cases (cell types or mouse genotypes). Normal distribution and equal variance was confirmed in the large majority of data and therefore, we assumed normality and equal variance for all samples. Based on this, we used the Student’s t-test (two-tailed, unpaired) to estimate statistical significance. For contingency tables, we used the Fisher’s exact test. Statistical analysis was performed using Prism (GraphPad Software, La Jolla, CA) or Microsoft Office Excel. All the experiments presented in this work were performed at least 3 times (between 3 and 10 independent biological replicates).

## Data availability

RNAseq data has been deposited in the GEO repository under accession number GSE202831.

## Acknowledgements

We thank the CNIO Histopathology, Molecular Imaging and Bioinformatics Units, UCM Confocal Unit, CNIO and UCM Animal Facilities for their technical support. We are indebted to the members of the Cannabinoid Signaling group (UCM) and the Cell Division and Cancer group (CNIO) for their constant support and advice. We are grateful to the “12 de Octubre” Hospital, particularly to the Radiology, Gynecology and Oncology Units for their constructive participation in this work and to the patients enrolled in this research study, who kindly donated one cylinder of their core-needle biopsy for the development of this project. This work has been in part financed by benefactors, through the crowdfunding project “Match point against breast cancer” (PRECIPITA PR242, 2019; FECYT; Spanish Ministry of Science and Innovation, MICINN, led by MS-R), and donations to “Asociación Española contra el Cáncer” (AECC). We are extremely thankful to all our donors. The work has been funded also by the Spanish Ministry of Economy and Competitiveness (supported with European Regional Development funds): PI20/00590 to CS. MS-R was supported by AECC (AIOA120833SALA and INVES18005SALA), a “Juan de la Cierva incorporación” and a “Ramón y Cajal” contract (from the MICINN). NGM-I was supported by AECC (PRDMA19003GARC).

## Author Contribution

NGM-I and MS-R performed most of the cellular and *in vivo* assays. SL and FM performed the micro-CTs and FM supervised the analysis of *in vivo* imaging data. PG and EJC performed the immunohistochemistry experiments and EJC supervised the IHC work. OG-C contributed to the analysis of the RNAseq data. JJM-O and AC-P helped with confocal imaging of organoids. MQ-F provided the MMTV-PyMT mouse model used in this study. EC, CS, SA, LS and SJ helped preparing the documents and protocols to be followed at the hospital for patient enrollment, recruited the patients, extracted the patient tissue samples and provided clinical advice. MS-R, CS and MM participated in project design, contributed to data evaluation and analysis and supervised the work. MS-R conceived the idea and wrote the manuscript, with the help of the main authors.

