## Supplemental Figures for "miR-203 drives breast cancer cell differentiation"

### (Supplementary Information)

Nuria G. Martínez-Illescas,[1,2,3] Silvia Leal,[4] Patricia González,[5] Osvaldo Graña-Castro,[6,7] Juan José Muñoz-Oliveira,[8] Alfonso Cortés-Peña,[8] Miguel Quintela-Fandino,[9] Eva Ciruelos,[10,2] Consuelo Sanz,[10,2] Sofía Aragón,[10,2] Leisy Sotolongo,[10,2] Sara Jiménez,[10,2] Eduardo J. Caleiras,[5] Francisca Mulero,[4] Cristina Sánchez,*[1,2] Marcos Malumbres,*[3] María Salazar-Roa*[1,2,3]

*The authors declare no competing financial interests.*

<u>Supplementary Information</u>

**Supplementary Figure S1.** PyMT mammary tumors at two different time points, treated or not with miR-203.

**Supplementary Figure S2.** miR-203-promoted morphological changes on mammary tumor organoids compared to those triggered by other well-known differentiation stimuli.

**Supplementary Figure S3.** RNA sequencing of organoid samples, derived from healthy or tumor tissue, and exposed *in vitro* to miR-203.

**Supplementary Figure S4.** Analysis of specific mRNA profiles for basal cells, development, cell migration, metabolism and cell cycle in organoid samples, derived from healthy or tumor tissue, and exposed *in vitro* to miR-203.

**Supplementary Table 1** (extended excel data for Supplementary Figure S4)

Supplementary Fig. 1

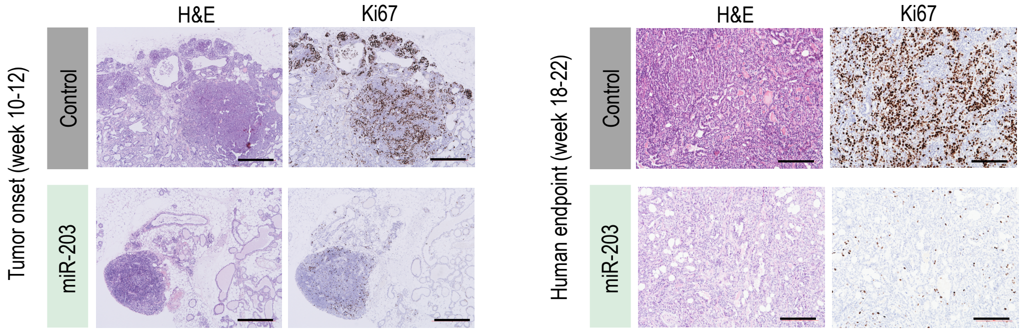


**Supplementary Fig. S1**. **PyMT mammary tumors at two different time points, treated or not with miR-203.** Illustrative immune-histopathological analysis of representative tumors (from miR-203 wild-type or miR-203 knock-in; PyMT mice exposed to Dox in vivo) and analyzed at two different time points: tumor onset (week 10-12, when tumors are not generally noticed by micro-CT yet, but identified by Hematoxylin and Eosin; left panel) or the experimental human endpoint (week 18-22; right panel). Hematoxylin and Eosin (H&E) and Ki-67 (as proliferation marker) staining are shown. Scale bar, 500 μm.

### Supplementary Fig. 2

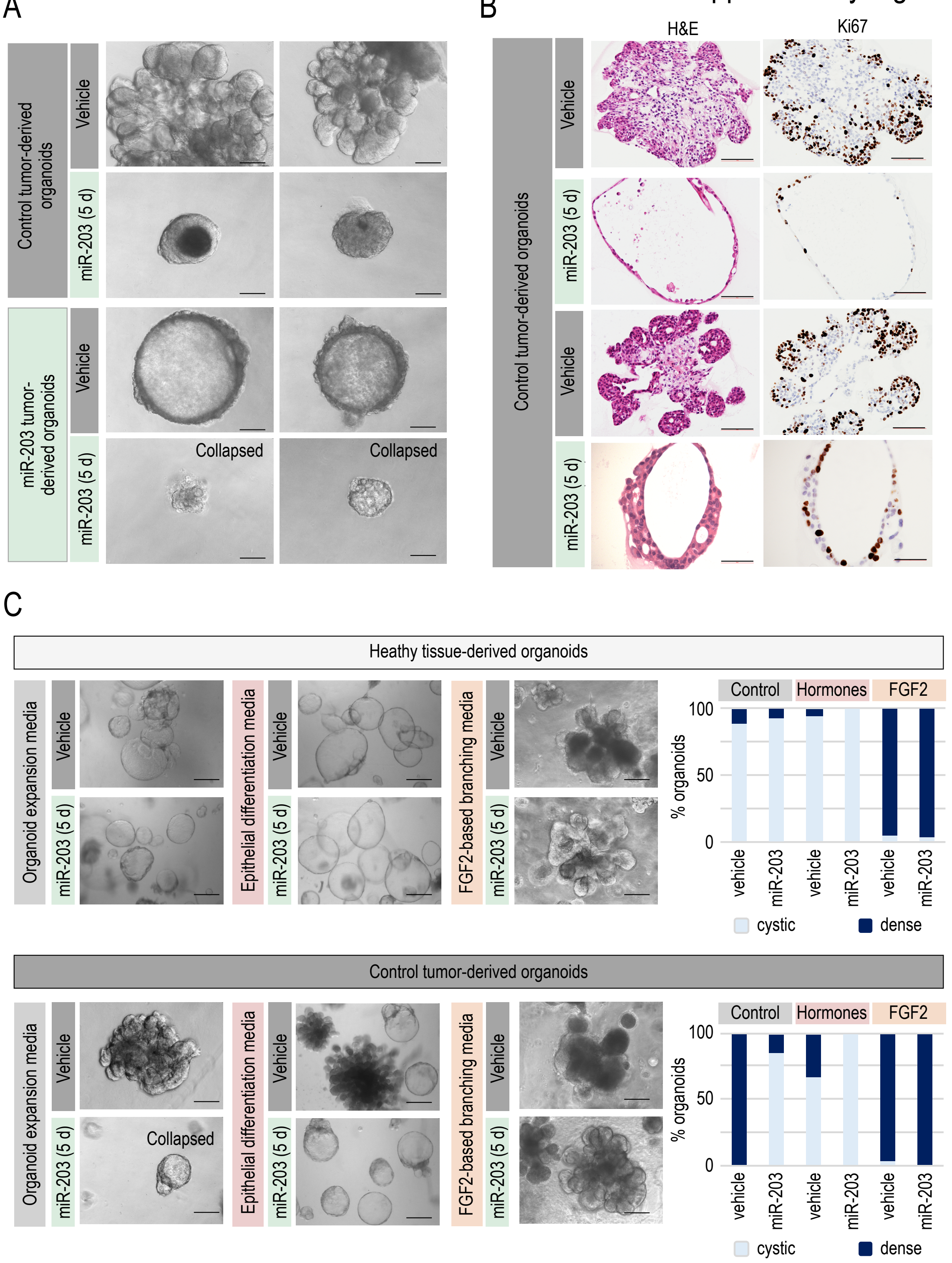

**Supplementary Fig. S2. miR-203-promoted morphological changes on mammary tumor organoids compared to those triggered by other well-known differentiation stimuli**. A, Representative bright-field images of tumor-derived organoids (tumors from miR-203 knock-in; PyMT mice treated in vivo either with vehicle or Dox), exposed in vitro to vehicle or miR-203 (Dox) during 5 days and followed by miR-203 withdrawal for 2 more weeks (indicated as “miR-203 5d” in the figure). Examples of collapsing organoids are shown, in the miR-203 exposed cultures. B, Representative examples of the immune-histopathological analysis of organoids generated and treated as in A. H&E and Ki-67 staining were tested. C, Representative bright-filed images of healthy tissue-derived organoids (upper panels) or control tumor-derived organoids (bottom panels), exposed in vitro during 2 weeks to the indicated treatments: left panels, organoids were cultured on basic expansion media, and treated with vehicle or miR-203 during the first 5 days; middle panels, organoids cultured either on epithelial differentiation media (consisting on prolactin, insulin, epidermal growth factor, hydrocortisone, bovine pituitary extract and gentamicin/amphotericin B), and treated with vehicle or miR-203 during the first 5 days; right panels, FGF2-based branching induction media, and treated with vehicle or miR-203 during the first 5 days. Quantification of the percentage of organoids exhibiting cystic or dense shapes respect to the total number of organoids is shown for each experimental setting and culture condition. Scale bar, 100 $\mu$m.

Supplementary Fig. 3

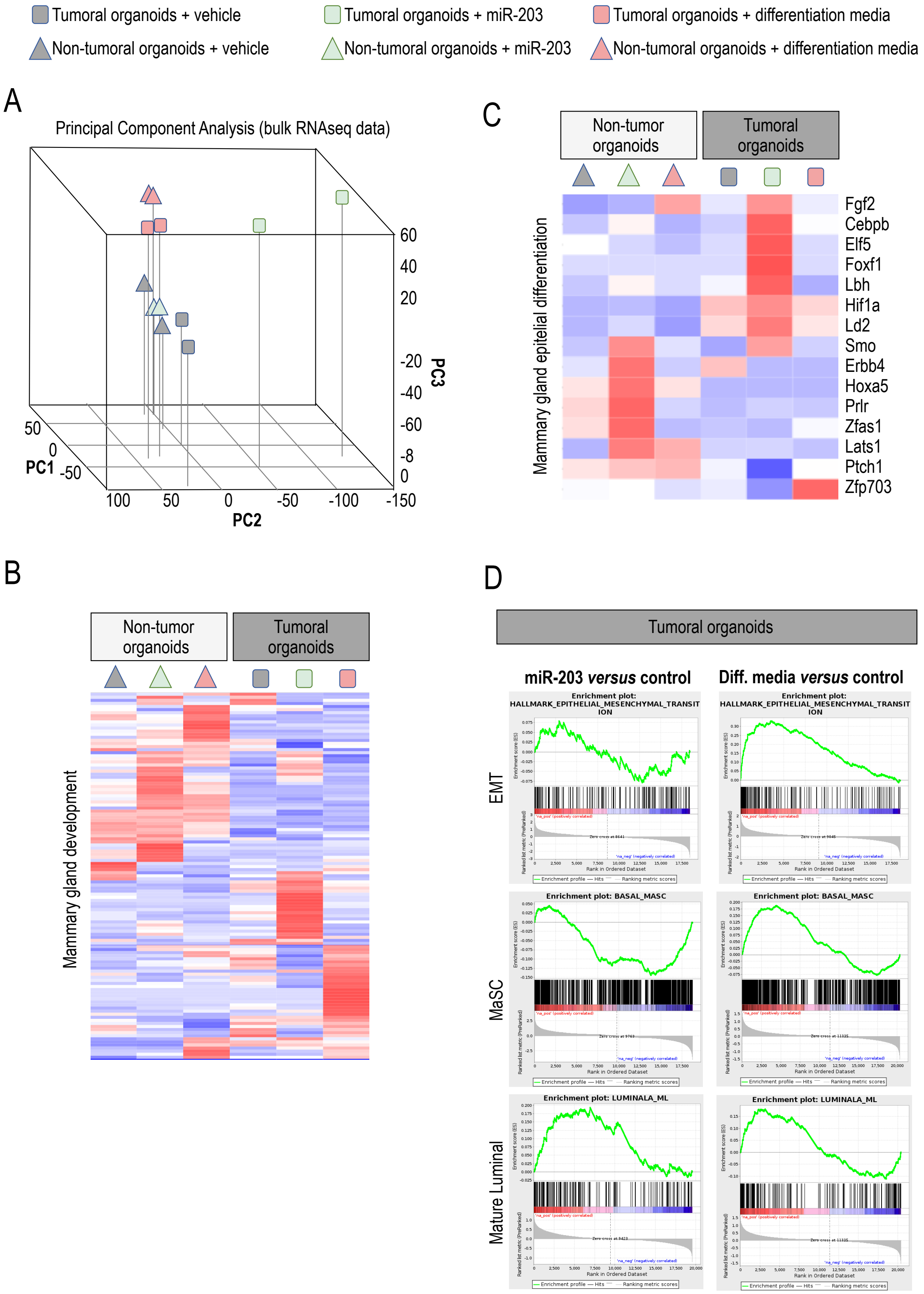

**Supplementary Fig. S3. RNA sequencing of organoid samples, derived from healthy or tumor tissue, and exposed *in vitro* to miR-203**. A, Principal Component Analysis of RNA sequencing performed on the samples indicated in the figure: non-tumor (triangles) or tumor (squares) mammary-gland-derived organoids, exposed to either vehicle treatment (in grey), miR-203 treatment for 5 days in vitro (in green) or differentiation media (consisting on prolactin, insulin, hEGF, hydrocortisone, BPE (bovine pituitary extract) and gentamicin/amphotericin B; in pink). The color and shapes code will be maintained throughout the figure, for clarity. B, C, Heatmaps showing the Gene Ontology signature for mammary gland development (B) and epithelial differentiation (C) in all the samples tested by RNA sequencing, as indicated (color and shape code as in A). D, Enrichment GSEA plots showing the induction of gene signatures characteristic of Epithelial-to-Mesenchymal Transition (EMT), Mammary Stem Cells (MaSC) and Mature Luminal cells in the comparisons indicated: miR-203-treated versus control tumor organoids (left panels) and differentiation media-cultured versus control tumor organoids (right panels).

### Supplementary Fig. 4

| Tumoral organoids | **miR-203 *versus* control: Significantly down-regulated genes** |
|---|---|

## A

**ARCHS4 Tissues data base**

| term | p-value | q-value | overlap_genes |
|---|---|---|---|
| Basal cells | 9.741974e-20 | 3.507111e-18 | FOXE1, FAM57A, GALNT18, TNC, FOXI1, CALML3, TFCP2L1, PRSS22, ESPN, DMKN, AQP3, WFDC5, PTPRF, NKPD1, HK2, GJA1, CYP26B1, NIPAL1, TRIM29, SLC16A3, KRT6A, BOK, KRTDAP, CERS3, ELOVL4, IL18, KRT5, OVOL1, OSMR, HSPG2, SERPINB8, PGF, EPN3, PROCR, SLPI, ELF3, ADGRF4, MALL, DDIT4, LY6D, FSCN1, KCTD11, DSG3, IVL, HBEGF, SEMA3B, IL20RB, LYPD3, SBSN, KLK8, CST6, NDRG1, BARX2, KLK7, KLK6, MTHFD1L, SH3BP1, STC2, P2RY1, PLEK2, SFN, LY6G6C, TSPAN1, NGFR, WNT10A, JUP, CAVIN3, KLK13, G0S2, EPHX3, KLF4, KLK10, KLK11, VEGFA, GJB2, KRT17, NLRP10, P4HA2, REEP4, KRT14, BCAR1 |

**PanglaoDB Augmented 2021**

| term | p-value | q-value | overlap_genes |
|---|---|---|---|
| Basal Cells | 9.941465e-14 | 4.473659e-12 | [BNIP3, KRT5, PRSS22, CST6, GJB2, SLPI, KRT17, TRIM29, ADGRF4, MALL, KRT14, LY6D, PLEK2, SFN, S100A14, SLC16A3, TSPAN1, KRT6A] |

## B

**Organ & Cell development**

| Term | GO | Count genes | % | P-value | Benjamini |
|---|---|---|---|---|---|
| Animal organ development | GO:0048513 | 71 | 30,5 | 2,8E-06 | 1,3E-03 |
| Cell development | GO:0048468 | 37 | 15,9 | 5,7E-02 | 7,2E-01 |

**Cell migration & Motility**

| | | | | | |
|---|---|---|---|---|---|
| Cell Migration | GO:0016477 | 33 | 14,2 | 1,3E-04 | 2,3E-02 |
| Regulation of cell migration | GO:0030334 | 24 | 10,3 | 2,8E-04 | 3,8E-02 |
| Regulation of cell motility | GO:2000145 | 24 | 10,3 | 6,4E-04 | 5,7E-02 |
| Positive regulation of cell migration | GO:0030335 | 15 | 6,4 | 3,6E-03 | 1,6E-01 |

**Metabolism**

| | | | | | |
|---|---|---|---|---|---|
| Protein metabolic process | GO:0019538 | 74 | 31,8 | 1,7E-03 | 1,1E-01 |
| Negative regulation of protein metabolic process | GO:0051248 | 26 | 11,2 | 1,4E-03 | 9,8E-02 |
| Regulation of protein metabolic process | GO:0051246 | 49 | 21,0 | 1,6E-03 | 1,0E-01 |
| negative regulation of cellular metabolic process | GO:0031324 | 46 | 19,7 | 2,3E-03 | 1,2E-01 |

**Cell cycle**

| | | | | | |
|---|---|---|---|---|---|
| Cell Cycle | GO:0007049 | 13 | 5,6 | 6,7E-02 | 1,0E+00 |

**Supplementary Fig. S4. Analysis of specific mRNA profiles for basal cells, development, cell migration, metabolism and cell cycle in organoid samples, derived from healthy or tumor tissue, and exposed *in vitro* to miR-203**. A, “Basal cell” signatures differentially down-regulated in miR-203-exposed tumor organoids versus their corresponding control counterparts, according to Enrichr (https://maayanlab.cloud/Enrichr/) (1) in two different data bases: ARCHS4 and PanglaoDB. B, Gene Ontology (GO) Signatures for “organ and cell development”, “cell migration and motility”, “metabolism” and “cell cycle” differentially down-regulated in miR-203-exposed tumor organoids versus their corresponding control counterparts, as defined by “The Database for Annotation, Visualization and Integrated Discovery” DAVID (https://david.ncifcrf.gov/) (2). All the genes recognized as significantly down-regulated in miR-203 treated versus control tumor samples and clustered in any of the signatures herein included are listed in the Figure and in the Supplementary Table 1.

References related to Supplementary Figure 4:

1. Xie Z, Bailey A, Kuleshov MV, Clarke DJB, Evangelista JE, Jenkins SL, et al. Gene Set Knowledge Discovery with Enrichr. Curr Protoc. 2021;1(3):e90.

2. Sherman BT, Hao M, Qiu J, Jiao X, Baseler MW, Lane HC, et al. DAVID: a web server for functional enrichment analysis and functional annotation of gene lists (2021 update). Nucleic Acids Res. 2022.
